# Segmentation and classification of retinal pigment granules in fluorescence lifetime imaging microscopy (FLIM) data

**DOI:** 10.64898/2026.06.29.735375

**Authors:** Maryam Ali, Hala Alhaj Ahmad, Hanan Alderzy, Martin Hammer, Rainer Heintzmann, Ondrej Stranik

## Abstract

Alterations of fluorescence properties in retinal pigment epithelium (RPE) cells caused by diseases such as age-related macular degeneration (AMD) highlight the need for detailed analysis of the fluorescent RPE granules at the individual level. Precise segmentation and classification of these granules remain challenging due to their limited visual separability. In this study, we present Classi4RPE, a computational algorithm designed to accurately segment RPE granules and classify them into three categories — lipofuscin (L), melanolipofuscin (ML), and melanin (M) — based on fluorescence lifetime imaging data, which provide distinctive contrast. The method is implemented in a custom Python framework and employs seeded watershed segmentation to isolate individual granules. Lipofuscin granules are identified as hyperfluorescent structures with longer lifetimes, while granules with shorter lifetimes are further analyzed based on their spatial lifetime distribution from the center to edge, enabling discrimination of ML from other melanin-rich granules. Our approach achieves high performance, with mean sensitivities of 0.99 for L granules and 0.90 for ML granules, and corresponding specificities of 0.93 and 0.98, respectively, compared to manually annotated ground truth. These results demonstrate the potential of Classi4RPE to surpass human visual limitations and provide a robust tool for quantitative RPE analysis.

## Introduction

Retinal Pigment Epithelium (RPE) granules are intracellular pigment accumulations formed from non-degradable phagocytic by-products. They are generated during the daily phagocytosis of photoreceptor outer segments as well as visual pigment recycling in the human retina **[1]**. Because the RPE serves as a critical metabolic and transport interface between the choroid and the neural retina **[2], [3]**, these granules contribute to structural and functional alterations associated with several retinal diseases [2], including age-related macular degeneration (AMD) **[2], [4]**. RPE granules are commonly categorized into three major types based on their autofluorescence properties and morphology. Lipofuscins (L) exhibit strong autofluorescence and typically display irregular, near-spherical shapes with heterogeneous fluorescence distributions. Melanin-rich granules, or melanosomes (M), show minimal autofluorescence in the visible range but stronger emission in the near-infrared (NIR) spectrum **[1], [5]**, and their morphology ranges from spherical to spindle-like. Melanolipofuscins (ML) contain both melanin and lipofuscin components; they characteristically exhibit a ring-like fluorescence pattern (melanin-rich core with fluorescent periphery) and may appear either spherical or spindle-shaped.

Fluorescence lifetime imaging microscopy (FLIM) provides an additional discriminative parameter that enhances granule characterization [6], and several recent studies have reported FLIM-based lifetime alterations associated with AMD progression [6], [7], [8], [9].

Quantifying these granules and understanding their statistical distributions is essential for assessing how biochemical alterations associated with retinal diseases influence RPE physiology. Advances in imaging and image-processing techniques have recently enabled more detailed clinical understanding of such granule-related changes [10]. Analyzing individual granules, thus, promises to achieve a deeper mechanistic understanding of correlations between granule types and AMD progression.

In this study, we used FLIM images acquired via two-photon excitation at 960 nm to perform automated segmentation and classification of RPE granules. This purpose requires higher resolved images and more efficient segmentation approach at the granule level, which is not typically addressed in most cellular-level studies. Conventional segmentation approaches previously applied to biological images [11] were unsuitable for this task because they required extensive manual fine-tuning, yielding a low reproducibility of the results. Similarly, recently developed software tools offering pretrained models for microscopic images [12], [13] still required manual adjustments for unresolved segments, which remained difficult to delineate due to the highly variable autofluorescence patterns within individual granules. It was also challenging and time consuming to create human-annotated training data, sufficient for training a neural-network based segmentation, thus limiting the feasibility of such neural-network approaches.

Previous studies have made substantial progress in segmenting retinal layers from optical coherence tomography images [14]. However, segmentation becomes considerably more challenging in the presence of RPE granules. Yu et al. (2023) applied semantic segmentation to RPE flat-mount images, successfully delineating RPE cells. Their findings highlight how AMD alters RPE cell borders, making it difficult to separate reliably during segmentation [15].

In contrast, our AMD-effected cross-sections of RPE images contain hundreds of granules belonging to multiple subclasses, each strongly affected by disease-related changes. This complexity necessitates instance-level segmentation to accurately delineate individual granules. To meet this need, seeded water shed segmentation [16] was applied by introducing seed assignment for each granule, enabling robust instance segmentation that captures nearly all granules present in the dataset. To further address these challenges, we developed a customized automated algorithm, Classi4RPE, capable of segmenting RPE granules and classifying them into three major granule types based on their FLIM characteristics.

## Methods

### RPE samples used

Use of human tissue was approved by institutional review board at the University of Alabama at Birmingham (#N170213002). Whole eyes of Caucasian donors aged 80 years or older without diabetes mellitus and no reported history of head trauma, surgeries affecting the retina, or conditions affecting the macula other than AMD were collected within 6 hours of death from Advancing Sight Network (Birmingham AL USA). After anterior segment removal, eyes were preserved in 4% buffered paraformaldehyde. Unstained glass slides with 12 µm thick cryosections were shipped on dry ice by overnight courier to Jena for autofluorescence lifetime microscopy (3-4 slides per eye).

### FLIM measurements

Autofluorescence lifetimes were recorded using an inverted multiphoton laser scanning microscope (Axio Observer Z.1 and LSM 710 NLO, 63x oil immersion objective Plan-Apochromat NA = 1.4, Carl Zeiss, Jena, Germany) in combination with a femtosecond Ti:Sapphire laser (Chameleon Ultra, Coherent Inc., Santa Clara, CA, repetition rate of 80 MHz with a pulse duration of 140 fs) and a single photon counting fluorescence lifetime imaging setup (Becker & Hickl GmbH, Berlin, Germany) with a 20 to 30 ps time resolution, as described [7]. The excitation wavelength for lifetime recordings was set to 960 nm for two-photon excitation (TPE). The lifetime imaging measurements are based on the principle of time-correlated single photon counting (TCSPC). The two-photon-excited fluorescence was recorded in two spectral channels. The short-wavelength spectral channel (SSC) was 500 – 550 nm, and the long-wavelength spectral channel (LSC) was 550 – 700 nm. All FLIM images were acquired as an average of 100 fast scans (mean photon count per pixel of approximately 1000, pixel dwell time of 3.1 µs) with a resolution of 256 × 256 pixel and a field of view of 27 × 27 µm.

The fluorescence decay images from the FLIM detectors were analysed using the software SPCImage 8.0 (Becker&Hickl GmbH, Berlin, Germany). For decay data fitting, a 3×3 pixel binning was applied. The decays where approximated by the fitting method of maximum likelihood estimation and with a three-exponential model yielding three decay time constants and three amplitudes.

### Structured illumination Microscopy (SIM)

As a technique with higher resolution than 2-photon FLIM, SIM was used to double-check for granule type. Images were taken using a commercial SIM device (ELYRA-S.1 system; Carl Zeiss, Jena, Germany) with a 40× NA 1.2 plan-apochromat (Carl Zeiss, Jena, Germany) water immersion objective lens combined with a 1.6× optovar. Excited with three different wavelengths 405nm (detected from 420 to 480nm and longer than 750nm), 488nm (detected from 495 to 550nm and longer than 750nm), 642 nm (detected from 655nm and longer). Presented images for this comprehension purpose were from Z-stack measurements taken with axial distance 3 or 4 µm with 21 slices and three rotations and 5 phase steps of the illumination pattern.

### Classi4RPE concept

Lifetime-fitted data were exported from the Becker & Hickl GmbH software together with the corresponding intensity images. The second detection channel (emission >550 nm) was selected for analysis, as it provided better signal quality for the approach described below; the discussion for this choice is addressed later. Figure 1 summarizes the workflow for RPE granules segmentation and classification.

**Fig 1.**
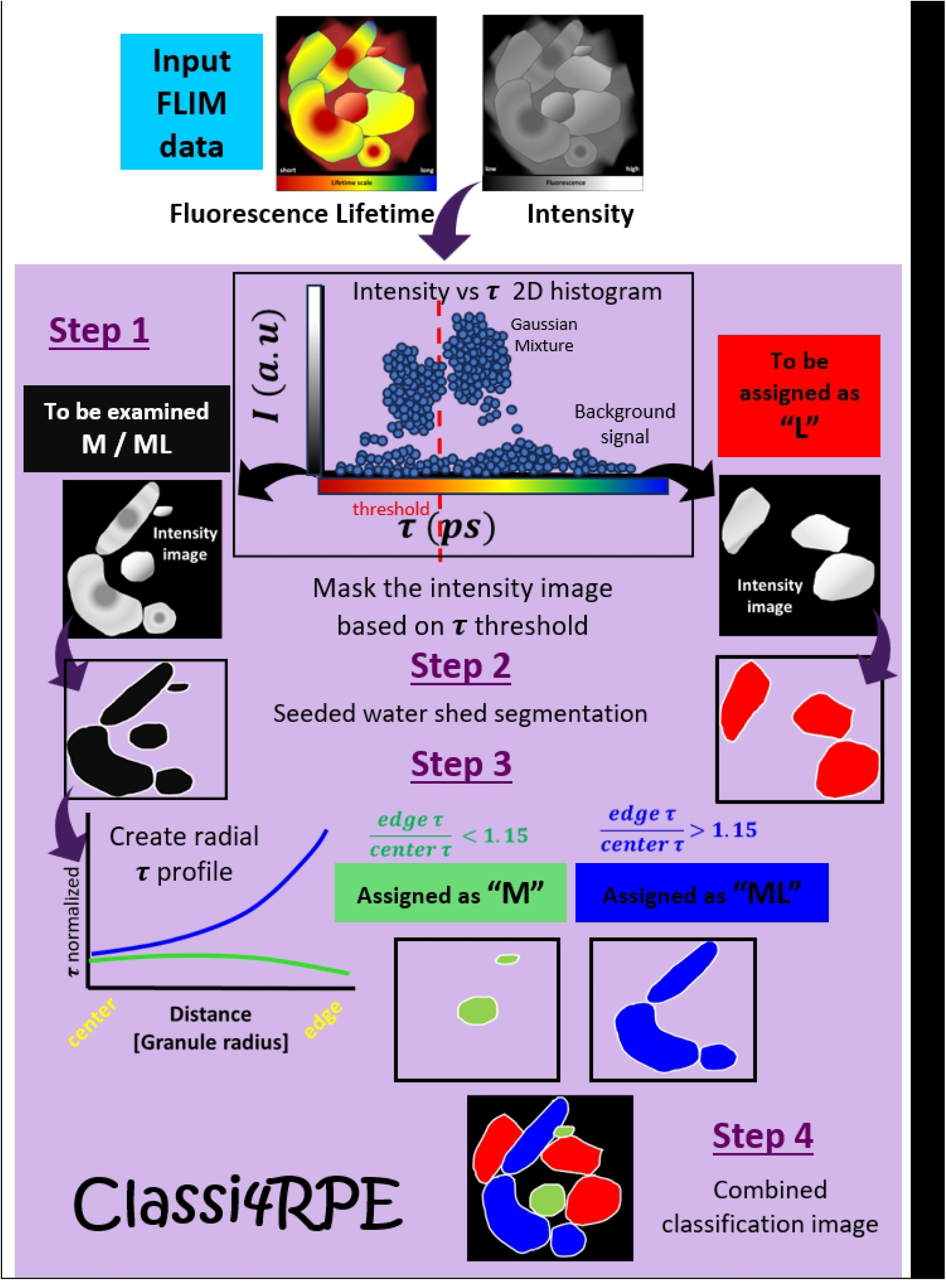
Classi4RPE workflow.

#### 1 Division of the image into two sub-images

The classification of granules is complicated by the heterogeneity of their fluorescence signal. Lipofuscins (L) exhibit characteristically high fluorescence intensity, whereas Melanosomes (M) produce a comparatively weak signal. The third granule type, Melanolipofuscin (ML), is a composite structure comprising a weak M-like signal in the center surrounded by the strong fluorescence characteristic of L. When a segmentation method (seeded watershed segmentation) is applied directly to the fluorescence intensity image, granules with weak signal (M and ML) are frequently under-detected, leading to erroneous classification outcomes. To overcome this limitation, we exploited additional information encoded in the spatial distribution of fluorescence lifetime, specifically the fact that L exhibits a longer fluorescence lifetime than M.

The fluorescence lifetime histogram (omitting the pixels without any significant intensity signal) was computed, and a threshold was determined as follows. The lifetime data were assumed to comprise three overlapping clusters, each described by a Gaussian distribution. A Gaussian mixture model (GMM) [17,18] was fitted to estimate the mean and standard deviation of each cluster (corresponding to M, L, and their intermediate state). The discriminative threshold was defined as the mean lifetime of the lowest-lifetime cluster (corresponding to M) plus 25% of its standard deviation. Pixels with lifetime values exceeding this threshold were assigned to Mask A, in which only L contributions are expected. The remaining pixels, with lifetime values below the threshold, were assigned to Mask B, which encompasses the M population and the central M-like region of ML granules, as described above.

#### 2 Segmentation of the two sub-images

Seeded watershed segmentation [16] was applied independently to each of the two masked intensity sub-images (Mask A and Mask B) to delineate individual granules. Seed points were identified as the local maxima of the difference-of-Gaussians (DoG, σ_1_,_2_= 1.6, 3.0) filtered intensity image. To prevent over-segmentation, a minimum inter-seed distance criterion was enforced: 7 pixels for Mask B (containing M and ML granules) and 3 pixels for Mask A (containing L granules), consistent with the given image magnification. A watershed algorithm was subsequently applied to the inverted intensity images, propagating segmentation regions outward from each seed point to delineate individual granule boundaries. Segmentation was performed using 8-connected neighborhood connectivity.

#### 3 Classification of the granules

Classification of L granules identified in the Mask A sub-image is straightforward. All cluster in the image are classified as L. Additionally; each segment is dilated by one pixel to ensure that granule edges are fully captured within the segmented region.

Classification of M and ML granules from the Mask B sub-image requires additional analysis. Segments are first dilated by two pixels to incorporate the outer ring of ML granules — a region dominated by L fluorescence and therefore absent from Mask B — into the subsequent analysis. For each dilated segment, a radial fluorescence lifetime profile is extracted and fitted with a quadratic function. For M granules, the radial lifetime distribution is spatially uniform. For ML granules, however, the radial profile exhibits systematically elevated lifetime values at the periphery relative to the center, reflecting the longer fluorescence lifetime of the L-rich outer shell compared to the M-like core. For each granule, the ratio of the lifetime at the edge to that at the center (R_τ) is computed. Granules with R_τ < 1.15 are classified as M, while those with R_τ ≥ 1.15 are classified as ML.

#### 4 Image merging (Classi4RPE output image)

The final Classi4RPE output image is reconstructed by combining all classified segments. As a consequence of the image subdivision and the dilation steps applied during segmentation, boundary overlap between adjacent granule segments may occur. In the final composite image, each granule type is rendered in a distinct color (L – red, ML – blue, M – green), and granule edges are highlighted in white to enable unambiguous distinction between adjacent granules of the same type.

Classi4RPE [19] is a custom-developed Python tool equipped with an interactive graphical user interface. It enables users to refine the classification and to export the resulting data, including the computed mean lifetime and associated segmentation information for each individual granule.

## Results

Classi4RPE was applied to seven datasets of which three were used to determine optimal threshold values, followed by evaluation of the overall performance across all datasets. Ground-truth images were generated from granule coordinates identified manually on the segmented images. However, not all granules could be clearly distinguished by eye, and in some cases the classification was particularly challenging. Consequently, segments that could not be reliably identified were excluded from the sensitivity and specificity analysis.

Across all datasets, the average sensitivity for detecting L granules was 0.99, and 0.90 for ML granules, with corresponding specificities of 0.93 and 0.98. M granules were excluded from the evaluation because some images did not contain them (based on manual inspection), and because distinguishing M from ML is inherently dependent on the imaging technique used (discussed later). Table 1 summarizes the statistical evaluation of the algorithm performance among the seven data sets, showing the sensitivity and specificity with and without considering the un-identified granules.

**Table 1.** statistical evaluation of sensitivity and specificity of ‘Classi4RPE’ performance compared to the manual ground truth.

|  |  |  | Including the not identified<br>by ground truth | Excluding the not identified<br>by ground truth |
| --- | --- | --- | --- | --- |
| **Lipofuscins (L)** | Sensitivity | Average | 0.99 | 0.99 |
|  |  | Minimum | 0.96 | 0.96 |
|  |  | Maximum | 1.0 | 1.0 |
|  | Specificity | Average | 0.39 | 0.93 |
|  |  | Minimum | 0.35 | 0.86 |
|  |  | Maximum | 0.45 | 1.0 |
| **Melanolipofuscins**<br>**(ML)** | Sensitivity | Average | 0.90 | 0.90 |
|  |  | Minimum | 0.78 | 0.78 |
|  |  | Maximum | 1.0 | 1.0 |
|  | Specificity | Average | 0.89 | 0.98 |
|  |  | Minimum | 0.80 | 0.97 |
|  |  | Maximum | 0.98 | 1.0 |

Figure 2 illustrates one of the stronger results obtained with Classi4RPE. 2a and 2b show the intensity and lifetime images from channel 2, with granule coordinates marked on the intensity image. Figure 2c displays the SIM image for the same position (used for validation), while 2d presents the Classi4RPE output and 2e shows the ground-truth classification, where L and ML granules are colored red and blue, respectively. White, unassigned granules were not labelled because they were either not clearly visible as granules or too ambiguous to classify. The accompanying table lists the sensitivity and specificity values calculated between the ground truth (2d) and the Classi4RPE output (2e), demonstrating the high classification performance, with nearly complete agreement. Both L and ML granules achieved a sensitivity and specificity of 1.0.

**Fig 2.**
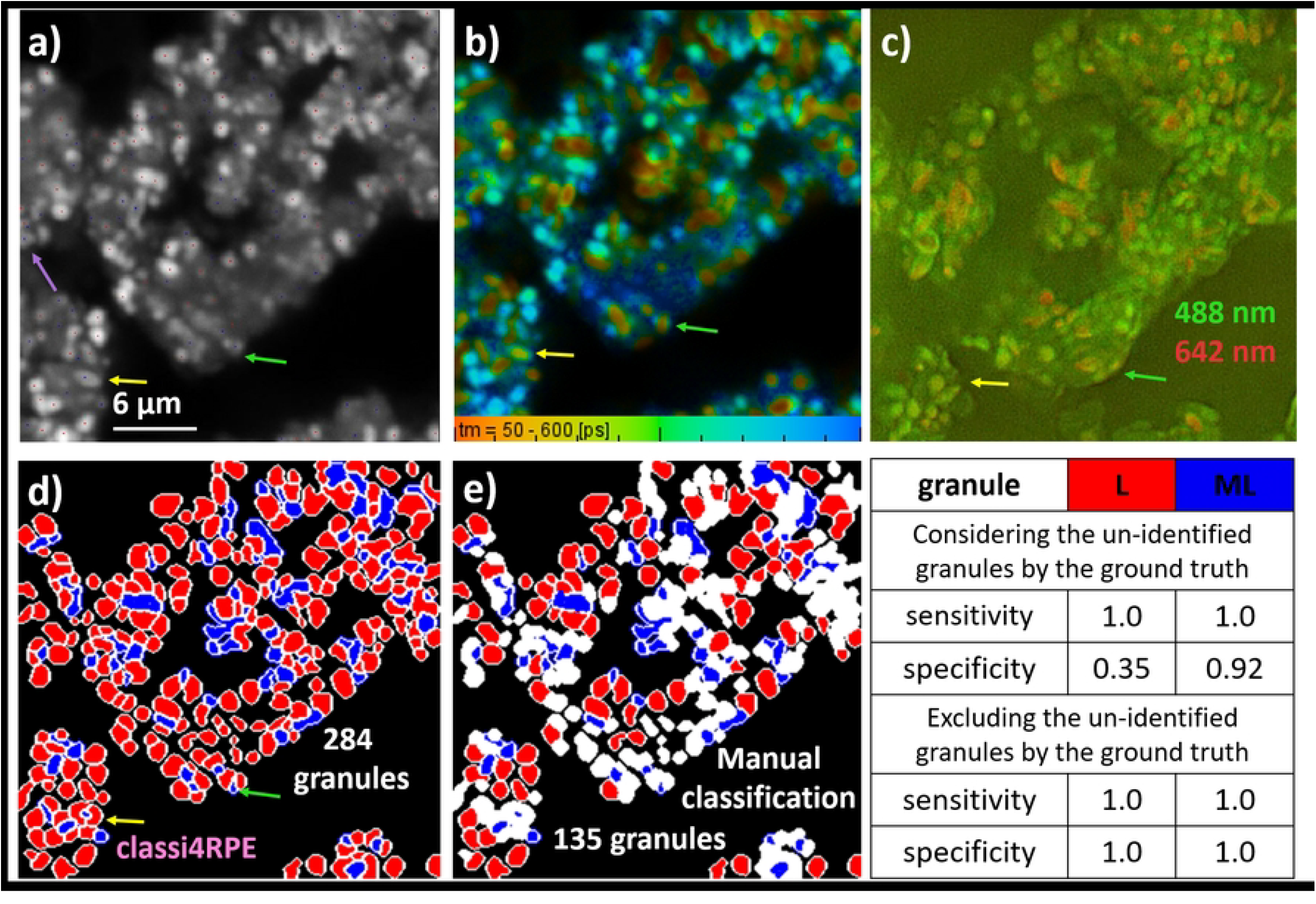
Classi4RPE high performance results of one of the tested FLIM data sets compared to the manual classification. a) 2-photon excitation (960 nm) intensity image detected by channel 2 (emission >555 nm). Marked dots referred to the manual assigned granules coordinates and classified as follows: red: L, blue: ML. b) corresponding FLIM image. c) SIM image for the same position measured with two wavelengths at 488nm (green) and 642nm (nm). d) Classi4RPE image. e) generated ground truth image from the manual classification with the same corresponding colors and the unidentified granules in white. e) Classi4RPE image. The provided table presents the calculated sensitivity and specificity for the results achieved in (e) compared to (d). Purple arrow: missed area in the segmentation. Yellow and green arrows: examples of sub-divided segmentation of one expected granule.

Regarding segmentation, the program detected 147 additional granules compared to manual annotation. Some regions were missed, such as the area indicated by the purple arrow. In addition, segmentation based on the intensity image influenced the identification of certain granules, which could represent large ML granules (yellow arrow) or clusters of granules that are visually difficult to resolve. Fortunately, these segmentation differences did not affect the classification outcome for this dataset.

In contrast, segmentation errors have an influence on the classification in the example shown in Figure 3. The yellow arrow highlights one ML granule that was only partially segmented, leading to its misclassification as L. The lifetime image in 3b and the SIM image in 3c both support its identity as ML, but the intensity-based segmentation caused the error. A similar issue occurred at the green arrow, where one ML granule was split into two MLs and one L. The purple arrow marks a complex region that was manually classified as a single large M granule; however, the intensity image revealed internal layers, causing the algorithm to segment it into multiple fragments classified as ML, which disagreed with the ground truth. The SIM image also shows internal compartments, making it difficult to determine whether the region should be considered M or ML. The orange arrow indicates another ambiguous granule that was classified as ML by the algorithm, although its identity remains unclear in the SIM image. The magnified region (orange box) reveals a surrounding ring-like structure in the lifetime image. However, this feature is difficult to interpret, as the observed border may originate from adjacent or underlying granules rather than the granule of interest. Due to this ambiguity, the sensitivity for ML and the specificity for L decreased to 0.98 in this dataset compared with the results shown in Figure 2.

**Fig 3.**
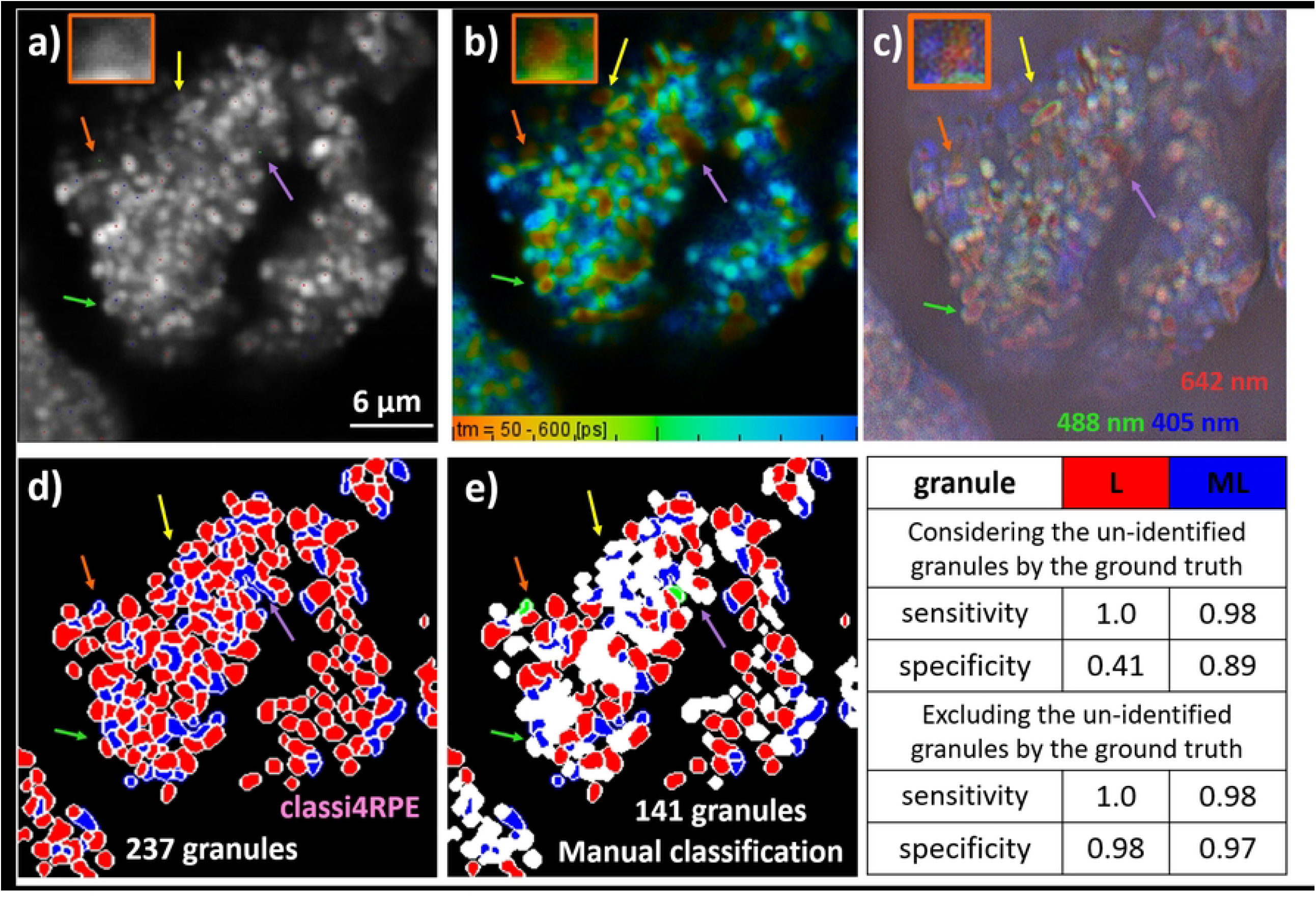
Classi4RPE results of another tested FLIM data sets compared to the manual classification. a) 2-photon excitation (960 nm) intensity image detected by channel 2 (emission >555 nm). Marked dots referred to the manual assigned granules coordinates and classified as follows: red: L, blue: ML, green: M. b) corresponding FLIM image. c) SIM image for the same position measured with three wavelengths at 488nm (green), 405 nm (blue), and 642nm (nm). d) Classi4RPE image. e) generated ground truth image from the manual classification with same corresponding colors and the unidentified granules in white. e) Classi4RPE image. The provided table presents the calculated sensitivity and specificity for the results achieved in (e) compared to (d). Purple, yellow and green arrows: examples of sub-divided segmentation of one expected granule that led to wrong classification. Orange arrow: unclear case of mis-classified Melanin granule (zoomed image in the orange box).

Figure 4 compares the results obtained from channel 1 (emission 500–550 nm) and channel 2 (emission >550 nm). 4a–c shows the intensity, lifetime, and Classi4RPE images for channel 1, while 4d–f shows the corresponding images for channel 2. The comparison demonstrates that segmentation and classification outcomes differ between channels. A slight increase in the number of segmented granules was observed in channel 2 compared to channel 1, resulting in broader coverage of both lipofuscin and short-lifetime granules (orange arrow). However, some granules were split into additional segments (yellow arrow), and differences in fluorescence between channels resulted in different total segment counts (4c vs. 4f). Furthermore, classification was affected both by segmentation differences and by variations in lifetime characteristics, resulting in a substantial increase in short-lifetime granules predominantly classified as M. This behavior appears unrealistic for certain granules. This figure illustrates why channel 2 was selected over channel 1, as the granules are more easily distinguishable, and the ring-like MLs are more clearly visible.

**Fig 4.**
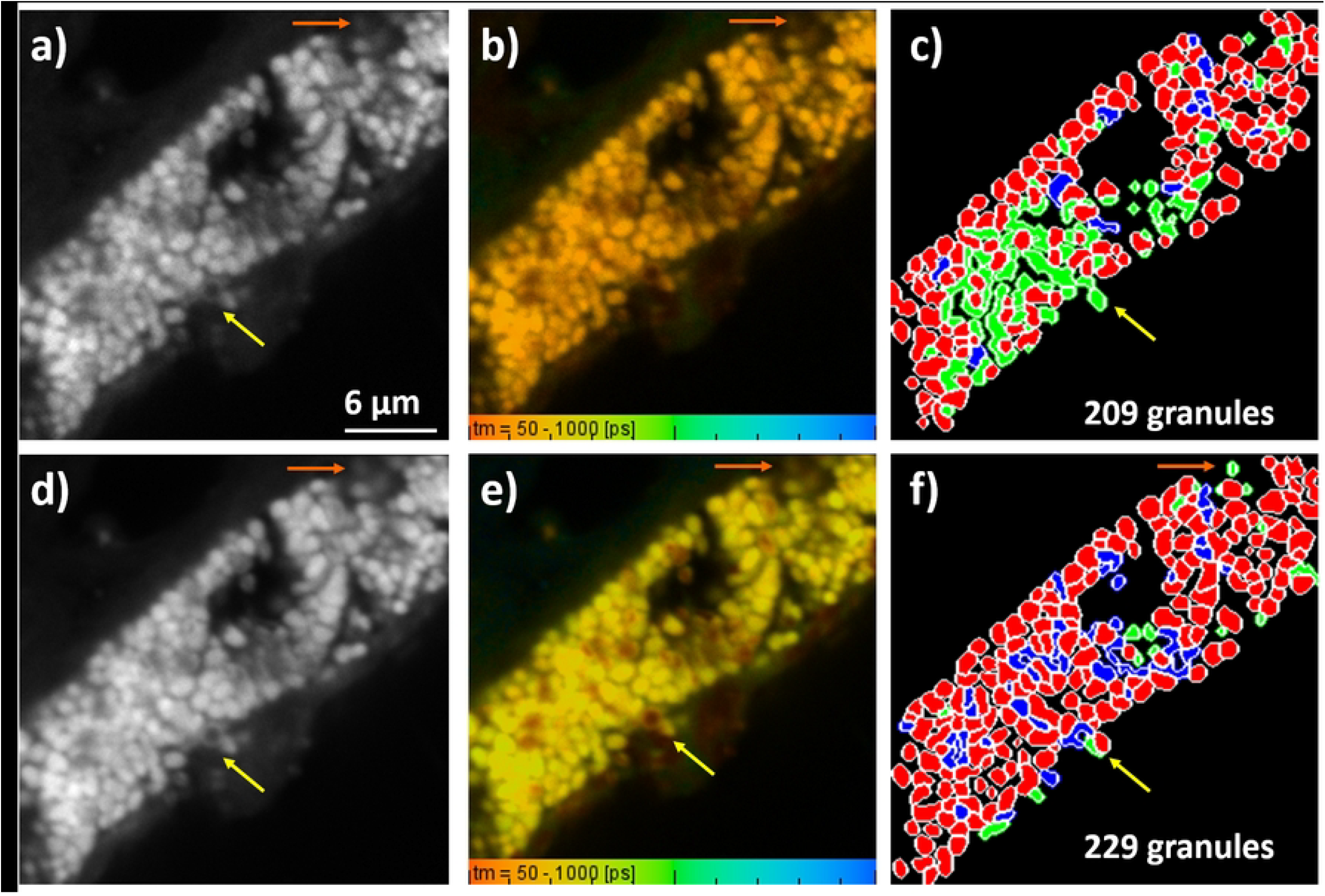
Classi4RPE results of channel 1 (emission 500–555 nm) and channel 2 (emission >550 nm). a-c) channel1. d-f) channel2. a and d) intensity images. b and e) corresponding FLIM images. c and f) Classi4RPE images with red: L, blue: ML, and green: M.

## Discussion

Classi4RPE, which demonstrates good sensitivity and specificity for L and ML granules, can successfully segment and classify RPE FLIM images. However, generating a reliable ground-truth image proved challenging. The morphology of RPE granules is highly complex [1], making it difficult even for human observers to distinguish individual granules or separate those that are clustered. Classification was particularly difficult for M versus ML granules, as it depends not only on lifetime-image contrast but also on the neighboring area. For example, when an ML or M granule is surrounded by L granules, delineating the boundaries between them becomes difficult. Similar ambiguity arises when granules overlap or appear stacked. Consequently, the ground truth included only granules that could be confidently identified by visual inspection, and excluding uncertain granules from sensitivity and specificity calculations was justified.

The highest sensitivity was achieved for detecting L granules, due to their distinct lifetime contrast, which applied as the first step in the classification pipeline (pre-classification). Nevertheless, achieving optimal segmentation of L granules remains challenging. Because segmentation was based on the intensity image, distinguishing true granules from sub-granular intensity variations was not always straightforward. This issue is particularly common for L granules, as previously reported by Bermond et al. (2020) [1]. Consequently, some L granules were over-segmented into multiple fragments—an outcome that cannot always be confirmed visually. These segmentation errors can lead to misclassification between L and ML granules, as illustrated in Figure 2: the yellow arrow indicates a large ML granule that was segmented into three parts, with the central segment classified as ML and the two outer segments as L. SIM imaging may also suggest the presence of multiple granules when viewed from a different imaging-modality perspective. The green arrow highlights a similar case for another granule.

These segmentation uncertainties can propagate into misclassification, as shown in Figure 3. The yellow arrow marks a ML granule that was incorrectly segmented into two granules (one classified as L and the other as ML) despite both FLIM and SIM images indicating that it is a single ML granule. A similar case is shown by the green arrow. One contributing factor may be the lifetime threshold used during pre-classification, which influences segmentation outcomes. Selecting this threshold is particularly challenging because RPE granules exhibit a wide range of lifetimes that vary between cells, especially in the case of pathologic cells migrated into the retina (examples in figs. 2 and 3), and the effects of AMD on these lifetimes remain poorly understood.

The purple arrow in Figure 3 highlights an especially complex case in which a large melanin-rich region appears as multiple layers in the intensity image and is therefore segmented into several granules. Although the lifetime image supports classification as M, SIM imaging does not clearly reveal how many granules are present or whether they should be labeled M or ML.

Distinguishing melanosomes (M) from ML granules remains difficult, contributing to the low sensitivity and specificity for the M class. Because the ground truth itself is uncertain for many M granules, we excluded M-related sensitivity and specificity from the evaluation. This challenge is illustrated by the orange arrow in Figure 3, where a granule classified as ML by the algorithm was labeled as M in the ground truth, yet SIM imaging also remains inconclusive. The side orange box on the left illustrates a magnified image on the same granule, and it shows the contrast between the borders (brighter) and the core (darker).

Another factor influencing M/ML discrimination is the ratio threshold between central and peripheral lifetime values. Although this threshold was optimized based on the available datasets, it requires a compromise between M and ML classification. Given our greater interest in accurately identifying ML granules, the threshold was tuned to favor ML detection, even at the expense of losing some M granules. In addition, the sensitivity and specificity for M were neglected because of the difficulty classifying them with certainty.

Finally, Classi4RPE, a newly developed algorithm incorporating characteristic lifetime profile computations of individual RPE granules for sub-classification, demonstrates strong performance in segmenting and classifying RPE into three types based on lifetime data, capturing details that are not discernible to the human eye. This is illustrated in Figures 2 and 3 (d, e), where the ground truth is compared with the Classi4RPE output. While segmentation performance is influenced by the quality of the input data, particularly with respect to detection channels, as shown in Figure 4, the algorithm remains robust across varying conditions Moreover, although the segmentation shows high sensitivity to the intensity image, especially in ML cases, where objects may be segmented separately due to thresholded lifetime pixels, the method still provides meaningful and consistent classification outcomes, even for challenging structures.

Further improvements could focus on optimizing lifetime thresholding and refining the segmentation criteria. In addition, the Classi4RPE framework may benefit from integration with higher-resolution imaging modalities or extension to three-dimensional analysis. Such developments could enable more accurate segmentation of SIM images and provide a more robust comparison with FLIM-based performance. Overall, despite the inherent challenges associated with the complex nature of the data, the algorithm achieves high performance. Its strong sensitivity and specificity relative to annotated ground truth— and its ability to capture features beyond visual interpretation—highlight its reliability as a tool for RPE granule analysis and its potential as a practical and effective method for studying RPE cells in AMD research.

## Conclusion

Age-related macular degeneration (AMD) induces significant alterations in RPE granules, necessitating further statistical investigation. Lifetime measurements provide important discriminatory information among these granules and can be used for their classification. However, visual classification remains challenging due to the difficulty of delineating granule boundaries, and some granules cannot be unambiguously assigned to a single type.

In this study, we developed a computational framework that provides a robust and efficient approach for RPE analysis, enabling systematic statistical tracking at the level of individual granules. This approach also mitigates the limitations associated with the lack of a reliable ground truth. The proposed algorithm, Classi4RPE, segments RPE images acquired via two-photon excitation using a seeded watershed approach and classifies granules into L, M, and ML types based on synchronized lifetime data.

High performance was achieved across seven datasets used to optimize the algorithm parameters, with almost complete coverage of the regions of interest and successful segmentation of most granules. This level of performance is difficult to achieve with conventional segmentation softwares, which often require extensive manual fine-tuning. Moreover, the algorithm demonstrated high sensitivity and specificity compared to manual ground truth annotations, highlighting its capability to exceed the limitations of human visual assessment in both segmentation and classification tasks.

Despite these promising results, several areas for improvement remain. Further refinement of segmentation parameters could enhance classification accuracy. Additionally, extending the framework to support three-dimensional SIM image segmentation represents a valuable direction for future work, potentially enabling higher-resolution analysis and improved structural characterization.

## Acknowledgments

Funding acknowledged from the German Research Foundation (DFG, Project: 542825796 “HiResi4RPE”).

## Data availability

Data used for testing can be obtained from the authors upon reasonable request. Python script used for “Classi4RPE” is available on GitHub: https://github.com/Maryam3Ali/Classi4RPE

